# Menin-MLL Inhibitor MI-503 Blocks Menin Nuclear Export and Suppresses Hypergastrinemia

**DOI:** 10.1101/2022.05.17.492246

**Authors:** Juanita L. Merchant, Zhen Wang, Sinju Sundaresan

## Abstract

Menin is the protein product of the Multiple Endocrine Neoplasia 1 (*MEN1*) gene locus at 11q13 and is a known tumor suppressor of neuroendocrine neoplasms (NENs). Gastrin-expressing NENs (gastrinomas) comprise the most frequent and malignant of the MEN1-dependent endocrine tumors. When gastrinomas are part of the MEN1 syndrome, they exhibit a greater propensity to develop within the submucosal Brunner’s glands of the duodenum. Therefore, models to analyze the biology of these intestinal gastrin-expressing NENs should consider their submucosal location.

**Aim:** The goal of this study was to determine whether the Menin-MLL inhibitor MI-503 suppressed hypergastrinemia.

**Methods:** A murine model of hypergastrinemia generated by omeprazole treatment of mice carrying a conditional deletion of *Men1* bred onto a somatostatin null genetic background (OMS) was treated intraperitoneally with MI-503 for 1 month. Primary enteric glial cells were prepared from these OMS mice and were treated with increasing doses of MI-503. Similarly human AGS and mouse STC-1 gastrin producing cell lines were treated with EGF without or with MI-503.

**>Results:** We found that the treatment reduced serum and gastro-duodenal tissue expression of gastrin. Ex vivo MI-503 treatment of glial fibrillary acidic protein (GFAP)+ enteric cells isolated from the OMS mice or gastrin-expressing cell lines revealed that MI-503 blocked the nuclear export of Menin and suppressed gastrin gene expression. RNA-Seq analysis of gastrin-treated GFAP+ enteric cells revealed that they express EGF receptor ligands and that EGF treatment of GFAP+ cells also induced Menin translocation and concurrent induction of gastrin gene expression.

**Conclusion:** We concluded that MI-503 inhibits gastrin gene expression by blocking Menin translocation.

## Introduction

Menin is the protein product of the Multiple Endocrine Neoplasia 1 (*MEN1*) gene locus at 11q13 and is a known tumor suppressor associated with neuroendocrine tumors (NETs) [1]. Gastrin-expressing neuroendocrine tumors (gastrinomas) comprise the most frequent and malignant of the *MEN1*-dependent endocrine cancers [2, 3]. Generally, sporadic gastrinomas occur in the head of the pancreas [2]. However, gastrinomas exhibit a greater propensity to develop within the submucosal Brunner’s glands of the duodenum that contribute to intestinal repair by producing mucins and EGF receptor ligands [4]. We generated a mouse model of hypergastrinemia generated by conditional deletion of *Men1* in enterocytes bred onto a somatostatin null (*Sst*^-/-^) genetic background and treated with omeprazole [5]. This Omeprazole-*Villin-Cre-Men1*^*FL/FL;*^ *Sst*^-/-^ (OMS) mouse model develops hyperplastic antral G cells [5]. In addition, the OMS mouse glial fibrillary acidic protein (GFAP)+ enteric glial cells express gastrin when nuclear menin translocates to the cytoplasm where it is ubiquitinated and degraded by the proteasome [5]. This novel mouse model exhibits several attributes observed in human subjects with *MEN1* gastrinomas, e.g., hypergastrinemia, submucosal gastrin-expressing cells in the duodenum and Type 2 gastric carcinoid tumors [5-7]. Moreover, duodenal gastrinomas (DGAST) from human subjects express neural crest markers, e.g., GFAP and S100b but not epithelial markers such as E-cadherin establishing the submucosal cells as enteric glia [5, 8].

Since >50% of duodenal gastrinomas do not exhibit loss of heterozygosity (LOH) [9, 10], it has been suggested that other mechanisms possibly initiate tumor development. The submucosal location of duodenal gastrinomas [4], lack of LOH identified in many neuroendocrine tumors carrying *MEN1* mutations [9, 10] and the absence of an in vivo model has hampered progress in developing new therapies. Clinical proteasome inhibitors such as bortezolmib are in clinical trials primarily for leukemia and have been shown to increase the stability of p21 and p27 [11, 12]. Ectopic expression of Menin mutants in the presence of the proteasome inhibitor MG132 blocked their degradation, but not their export to the cytoplasm [13, 14]. Small molecules that specifically prevent the nuclear export of Menin and its subsequent degradation in the cytoplasm, in contrast to simply inhibiting the proteasome might be an alternative mechanism to increase nuclear Menin levels. Menin-MLL (Mixed Lineage Leukemia) inhibitors (MIs) are small molecules that bind directly to menin at residues F9 and P13 [15, 16] where both JunD and MLL dock. With respect to leukemias, menin acts as a co-oncoprotein by directing mutated MLL to pro-proliferative target genes [16, 17]. Although the interaction of MLL protein with menin in the F9-P13 “pocket” [17] has been the most intensively studied mechanism [18], Menin binds several other proteins including GFAP [19, 20]. The transforming capability of MLL has also been examined in prostate cancer where MIs were shown to block proliferation of this solid tumor [17]. In addition, the authors found that Menin protein levels increase with MI treatment [17].

Since the OMS mice exhibited hypergastrinemia in the setting of non-cell autonomous loss of nuclear Menin in the enteric glial cells [5], we treated these mice with MI-503 and found that it blocks the nuclear export of menin resulting in suppression of plasma gastrin these mice.

## Results

### MI-503 blocks nuclear export of menin and suppresses hypergastrinemia

Menin-MLL inhibitors (MIs) are cyano-indole compounds that bind directly to Menin and facilitate its stability to heat [17]. We considered that this property of MIs might correlate with Menin stabilization. To test this possibility, we examined whether MI-503 was able to suppress hypergastrinemia in the *OMS* mice [5]. Three doses (10, 30, 55 mg/kg) of MI-503 were administered to the hypergastrinemic *OMS* mice by daily intraperitoneal injections for 4 weeks. There was a dose-dependent decrease in plasma gastrin with a maximum 70% decrease observed with the 55 mg/kg dose (**Fig. 1A**). Macroscopically there was a reduction in the size of the hyperplastic corpus, consistent with reduced gastrin levels (**Fig. 1B**). Menin protein increased in the villus cores and coincided with reduced gastrin peptide (**Fig. 1C,D**). Isolating enteric glial cells from the MI-503-treated *OMS* mice permitted determination of the gastrin content and demonstrated a significant 70% reduction in the peptide at the 55 mg/kg dose (**Fig. 1D,E**). A dose-dependent re-expression of menin was also observed in the glial cultures and coincided with reduced gastrin expression (**Fig. 1E**). Menin was retained in the nuclear fractions and was barely detectable in the cytoplasmic fractions of glial cells isolated from MI-503 treated *OMS* mice, compared to that of vehicle treated controls (**Fig. 1E**). Changes in nuclear versus cytoplasmic Menin were quantified by western blot (**Fig. 1F,G**).

**Figure 1.**
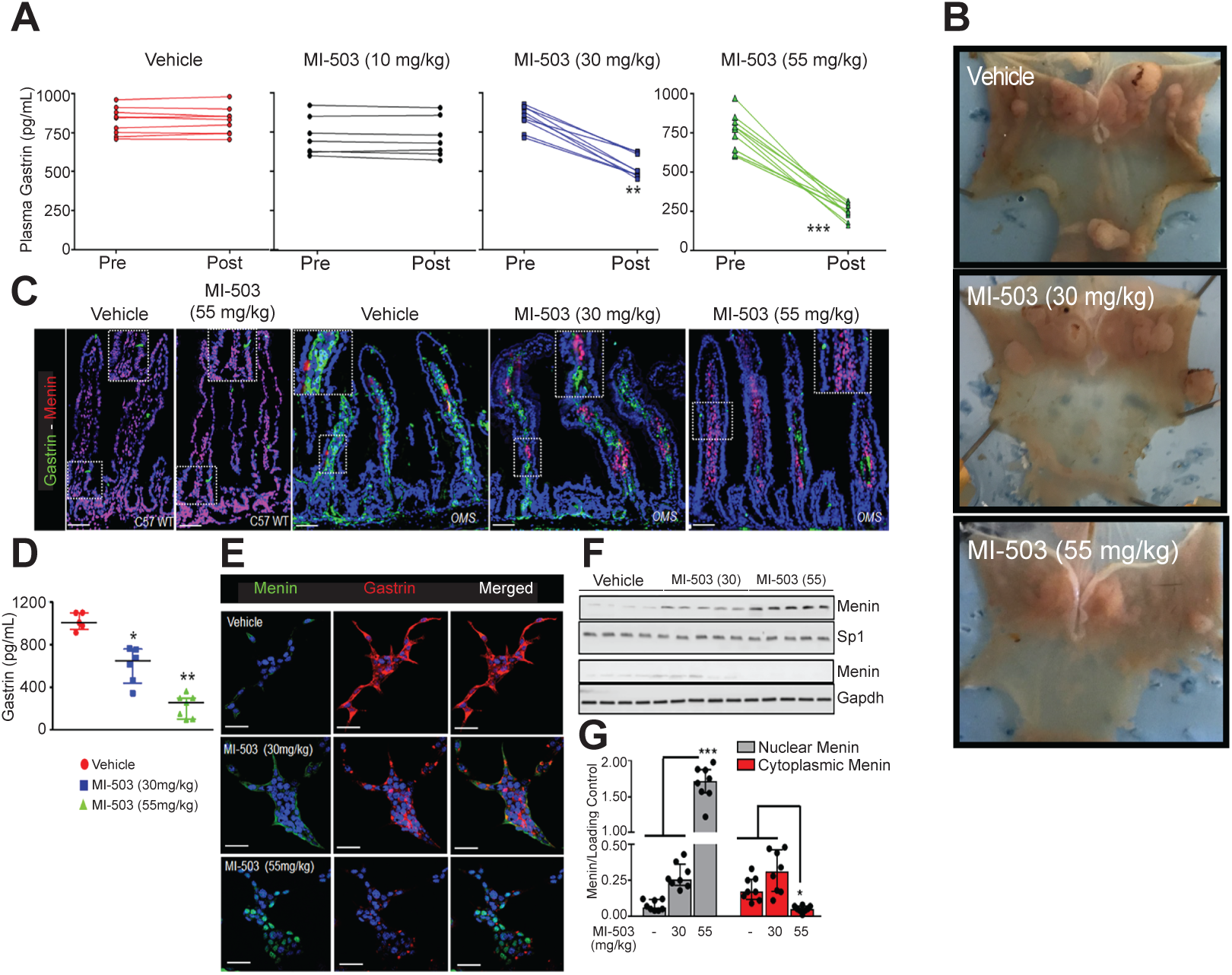
MI-503 suppresses hypergastrinemia and blocks nuclear export of Menin. (**A**) EIA measurement of plasma gastrin concentration from *OMS* mice before (Pre) and after (Post) treatment with vehicle, 10 mg/kg, 30 mg/kg, or 55 mg/kg of MI-503 for 4 weeks (*n=* 10-12 mice). (**B**) Representative photos of the stomachs from OMS mice treated with vehicle or the highest doses of MI-503, i.e., 30 or 55 mg/kg. Dose-dependent stabilization of Menin by MI-503. (**C**) Immunofluorescent staining of Menin (red) and gastrin (green) from duodenums of *OMS* mice after treatment with vehicle, 30 mg/kg, or 55 mg/kg MI-503 for 4 weeks. (**D**) EIA measurement of intracellular gastrin content of glial cultures isolated from omeprazole treated *Men1*^Δ*IEC*^*;Sst*^*-/-*^ mice after treatment with vehicle, 30 mg/kg, or 55 mg/kg MI-503 for 4 weeks (*n=*6-8 mice), expressed after normalization to total cellular protein content. (**E**) Immunofluorescent staining of gastrin (red) and menin (green) in glial cultures isolated from duodenal lamina propria of omeprazole treated *Men1*^Δ*IEC*^*;Sst*^*-/-*^ mice and then treated *ex vivo* with vehicle, 0.1 nM, or 5 nM MI-503 for 2 days. (**F**) Representative western blot showing Menin expression in total cell lysates from duodenal glial cultures (from *OMS* mice) treated with MI-503 for 2 days (N=4-5 mice) (**G**) Quantitation of Menin protein levels on blots as a function of MI-503 dose. (N= 8 mice)

To whether determine whether MI-503 affected nuclear export or degradation, we treated enteric glial cultures from the *OMS* mice with increasing amounts of MI-503 and found that 5 nM was sufficient to retain Menin in the nucleus *ex vivo* (**Fig. 2A-C**). If enteric glial cells from WT mice were treated with gastrin to induce Menin export as previously reported, MI-503 blocked the nuclear export as observed with leptomycin B (LMB) (**Fig. 2D,E**) coincident with dose-dependent reduction of ubiquitination and Menin degradation (**Fig. 2F,G**). Thus, we concluded that MI-503 blocks the nuclear export of Menin in vivo, which subsequently prevents menin ubiquitination and degradation in the cytoplasm.

**Figure 2.**
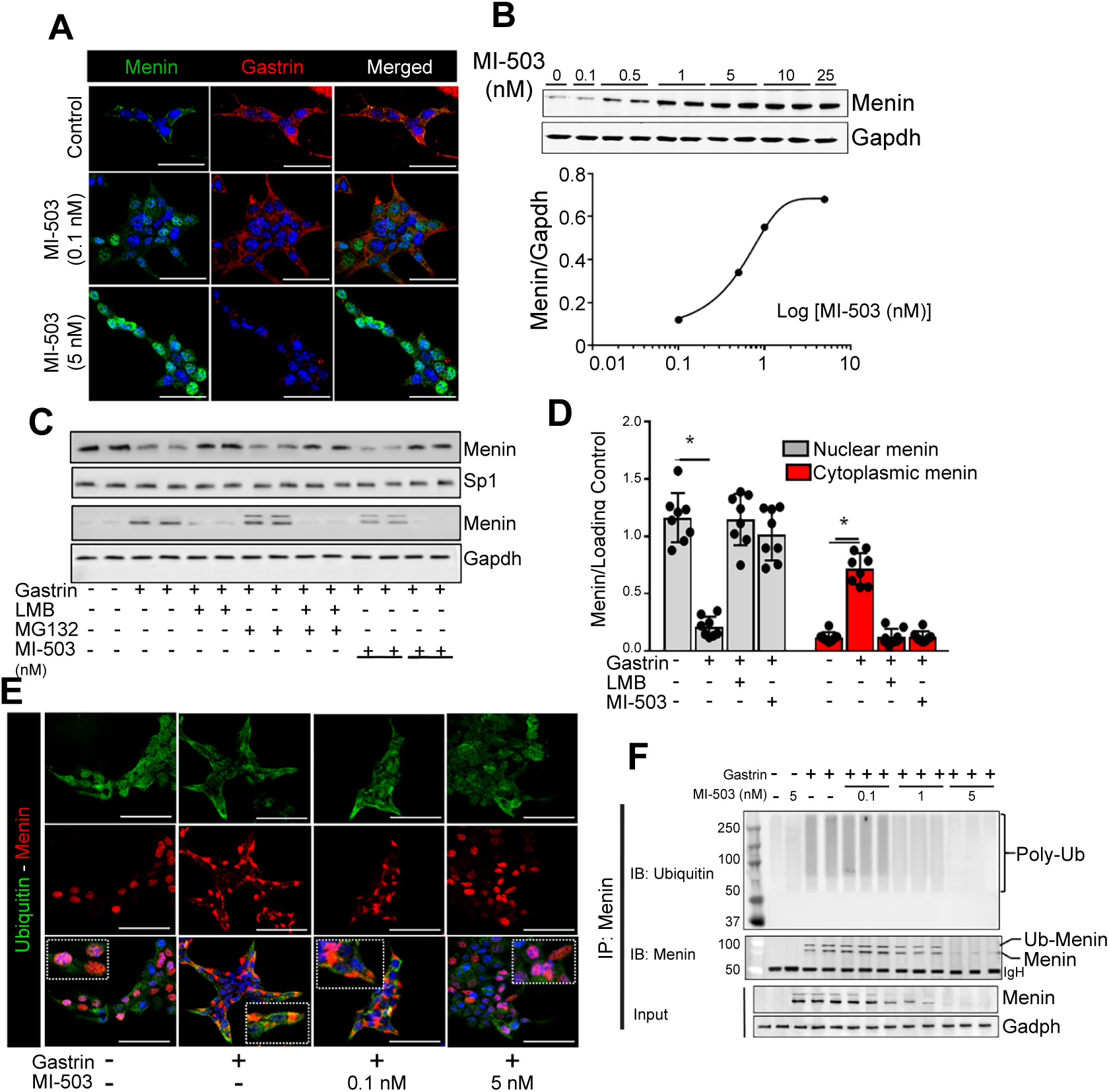
MI-503 blocks menin nuclear export and gastrin expression. (**A**) Dose-dependent stabilization of Menin observed after treating duodenal glial cultures prepared from OMS mice with 0.1 or 5 nM MI-503 for 2 days (Western blot for Menin; GAPDH loading control). (**B**) Menin/GAPDH ratio plotted as a function of MI-503 concentration; (**C**) Representative blot showing Menin expression with or without gastrin treatment of enteric glial cells for 2 days. Effect of 10 µm Leptomycin b (LMB) or MI-503 treatment. The proteasome inhibitor MG132 added to block degradation of ubiquitinated Menin; (**D**) Quantitation of Menin expression in nuclear and cytoplasmic fractions of glial cultures (from *OMS* mice) treated with or without gastrin in the presence or absence of Leptomycin B (LMB,10 µM), or MI-503 (5 nM) for 2 days. Data are expressed as integrated band intensities normalized to appropriate loading controls (Sp1 for nuclear fraction; GAPDH for cytoplasm) (*n*=8 expts). (**E**) Immunofluorescent staining for Menin (red) and ubiquitin (green) in glial cultures from C57BL/6 WT mice treated without or with 20 nm gastrin and ±MI-503 for up to 48 hours.

To dissect the role of gastrin in regulating glial cell Menin, we performed RNA-Seq on primary mouse glial cultures treated with gastrin. The volcano plot of genes induced demonstrated an increase in two EGFR ligands—epiregulin (Ereg) and neuregulin 1 (NRG1) (**Fig. 3A**). Quantitative PCR confirmed the increase Ereg and NRG1 mRNA in glial cultures (**Fig. 3B**). These results suggested that gastrin might also induce the release of EGF receptor ligands that subsequently induce menin translocation. Since the in vivo treatments used omeprazole (OM), we demonstrated that adding OM did not affect Ereg or Nrg1 mRNA (**Fig. 3C**). Next, we treated the enteric glial cultures with EGF and the receptor antagonist erlotinib and showed that erlotinib prevented gastrin-induced menin translocation to the cytoplasm (**Fig. 3D**). Therefore, we concluded that gastrin induces the expression of EGF receptor ligands that in turn can also modulate menin translocation. Therefore, gastrin could also regulate menin stability through the release of EGF receptor ligands.

**Figure 3:**
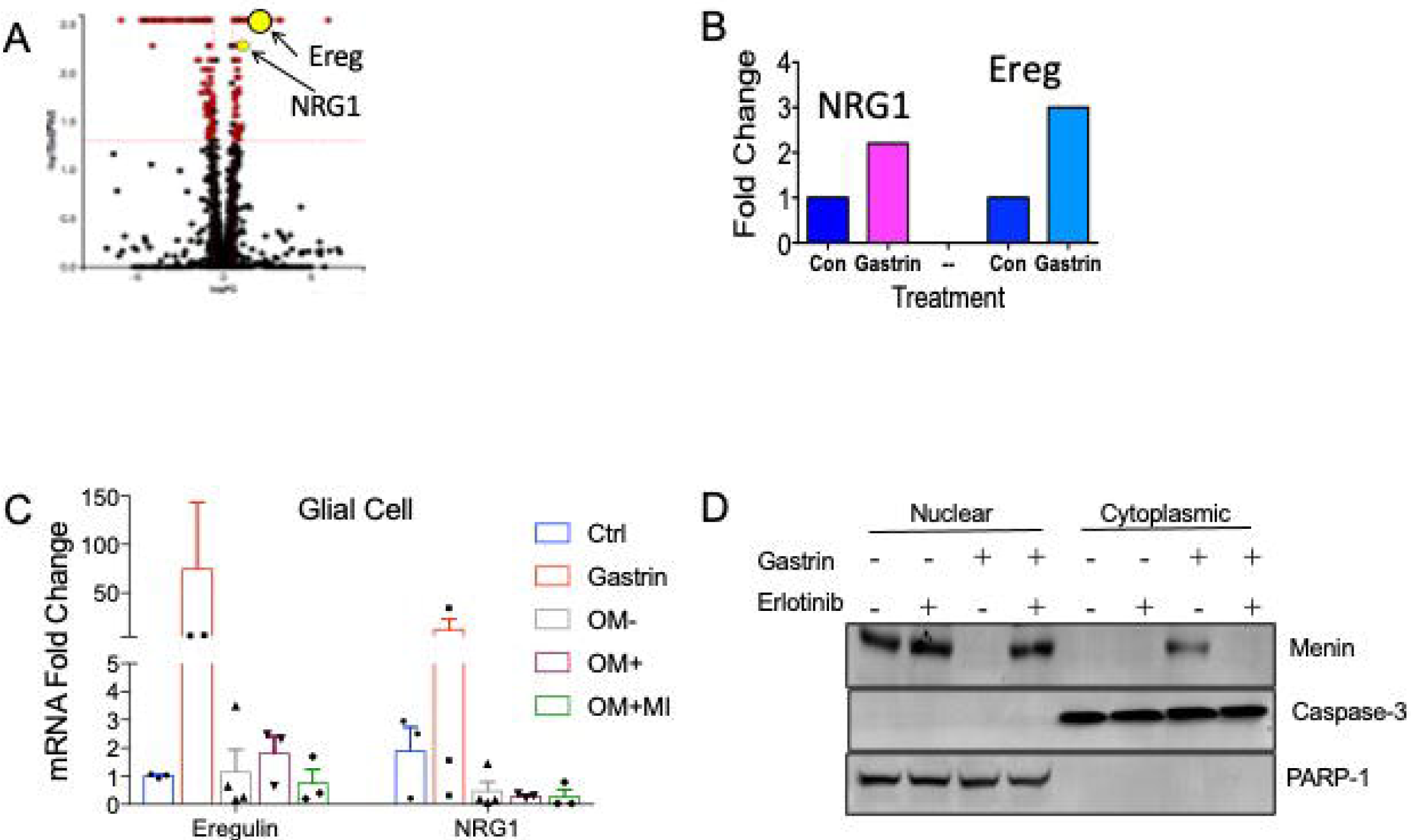
RNA-Seq of gastrin treated primary glial cells. **(A**) Volcano plot of upregulated and down-regulated genes expressed by enteric glial cells treated with gastrin; (**B**) Levels from RNA-Seq plotted as a function of treatment; (**C**) Levels of epiregulin or Neuregulin as a function of gastrin or MI-503 treatment; (**D**) Western blot showing effect of erlotinib (EGF receptor antagonist) on glial cells treated with gastrin.

We previously reported that activation of the EGF receptor is a potent inducer of gastrin gene expression [21]. Moreover, menin strongly suppresses gastrin gene expression in vitro and in vivo [5, 22, 23]. Somatostatin is a known inhibitor of gastrin expression and secretion exerts its effect in part by inhibiting protein kinase A and increasing menin expression [5, 24]. However, it is not known whether EGFR regulation of gastrin modulates menin protein. Since gastrin through the CCKBR receptor and PKA stimulates the nuclear export of menin in enteric glial cells that in turn de-represses gastrin expression [5], we tested whether EGF also induced the export of Menin and subsequently gastrin expression. Directly treating primary glial cultures isolated from C57BL/6 (wild type) mice with EGF for 16h, we found dose-dependent export of nuclear Menin, which subsequently induced gastrin expression (**Fig. 4A-D**). At least 10 nM EGF was sufficient to induce the nuclear translocation of Menin determined by a western blot of the nuclear and cytoplasmic fractions (**Fig. 4E**). PARP-1 and caspase-3 were used to confirm separation of the nuclear and cytoplasmic fractions occurred within 8 h (**Fig. 4E,F**). By 24h, there was a decrease in cytoplasmic menin indicative of its degradation by the proteasome (**Fig. 4E,F**). Treating glial cultures with bombesin 48 h after EGF treatment induced the release of gastrin peptide into the media (**Fig. 4G**). Therefore, EGF was sufficient to induce the nuclear export and degradation of menin.

**Figure 4:**
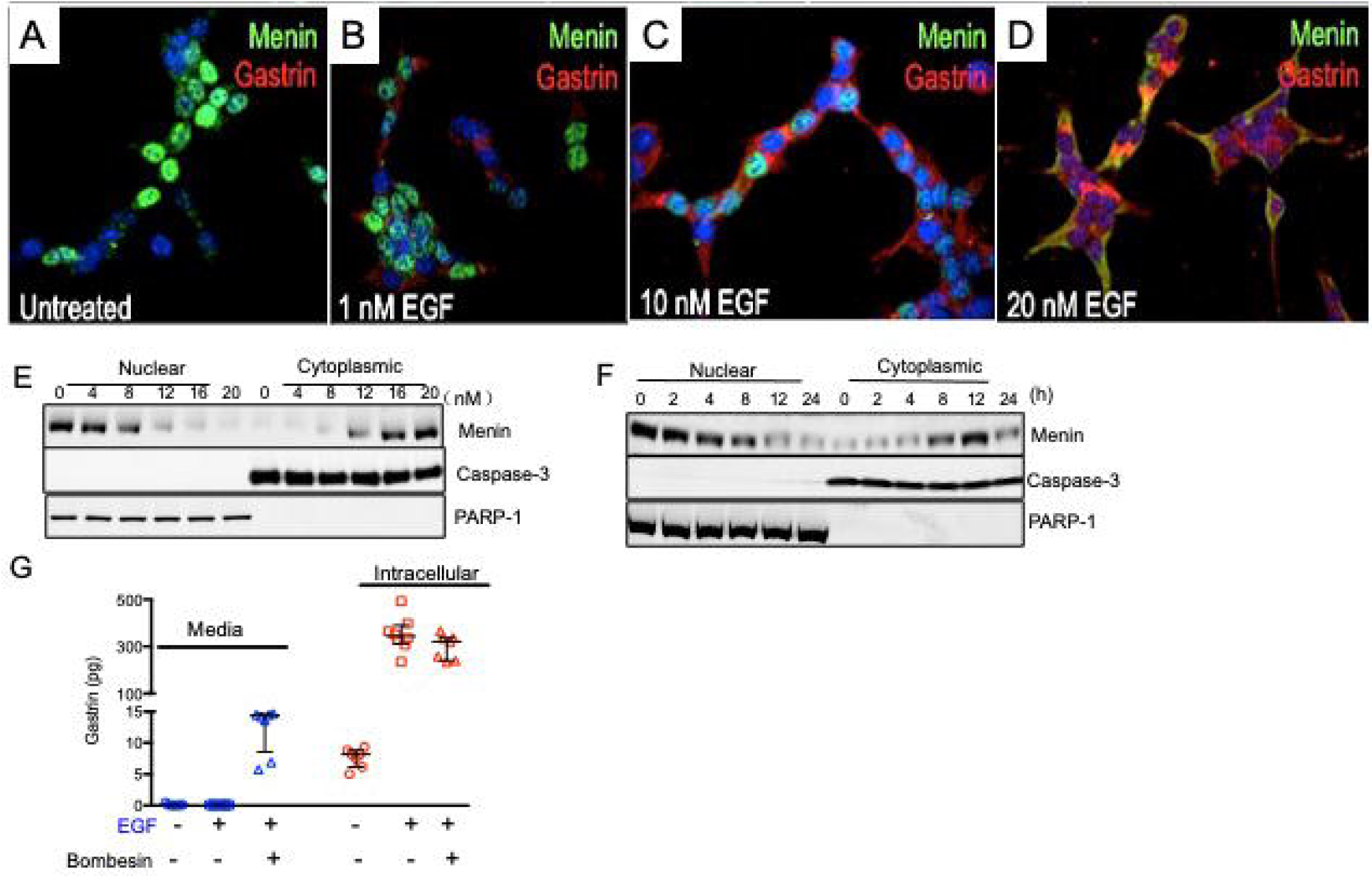
EGF induces gastrin gene expression through menin nuclear export and degradation. **(A)** Immunofluorescent staining for menin (green) and gastrin (red) after 0 to 20 nM EGF treatment of enteric glial cells. **(B)** Dose-dependent changes in the subcellular localization of menin with EGF at 12h. (**C)** Time dependent changes in the subcellular localization of menin with 20 nM EGF from 0 to 24h. After 48 to 72 h in culture, the enteric glial cell cultures the media was replaced with serum free media for 4h prior to EGF treatment. Nuclear and cytoplasmic fractions were generated and resolved by SDS-PAGE. Shown are the representative changes for N=3 experiments using PARP-1 and caspase-3 to document nuclear and cytoplasmic fractions respectively. (**D**) Quantitation for the time course shown in (**C**). (**E**) qPCR for menin and gastrin mRNA from the time course of EGF treatment of enteric glial cultures over 24h with 20 nM EGF. (**F**) Intracellular gastrin concentration determined by ELISA in glial cultures treated with 20 nM EGF. (**G**) Bombesin-induced secretion of gastrin from glial cultures after 72h in culture (N= 7 glial preparations). Data shown are the mean ± SEM. Kruskal-Wallis ANOVA with Dunn’s test of multiple comparisons to determine significance. * p< 0.05.

### Effect of MI-503 on cell growth

To determine if MI-503 affects cell growth in addition to blocking menin translocation, we treated three cell lines with MI-503, the human gastric adenocarcinoma AGS line, AGS cells stably overexpressing the gastrin receptor CCKBR (AGS-E), and the murine small intestinal enteroendocrine STC-1 cell line. MI-503 inhibited cell growth after 5 days of treatment with >1nM (**Fig. 5A**). To test whether MI-503 affected menin or gastrin gene expression AGS cells were treated with 20 nM gastrin and 10 nM MI-503 for up to 48 h (**Fig. 5B-D**). Quantitation showed that the decrease in Menin protein in this human cell line was greatest at 48h and was reversed with MI-503 treatment (**Fig. 5B,C**). Although MI-503 had no effect on *Men1* mRNA (**Fig. 5E**), MI-503 inhibited induction of gastrin gene expression (**Fig. 5F**) and restored nuclear Menin (**Fig. 5D**). Collectively, these results demonstrated that MI-503 blocked hypergastrinemia preventing the nuclear export of Menin, CCKBR induction and PKA activation.

**Figure 5.**
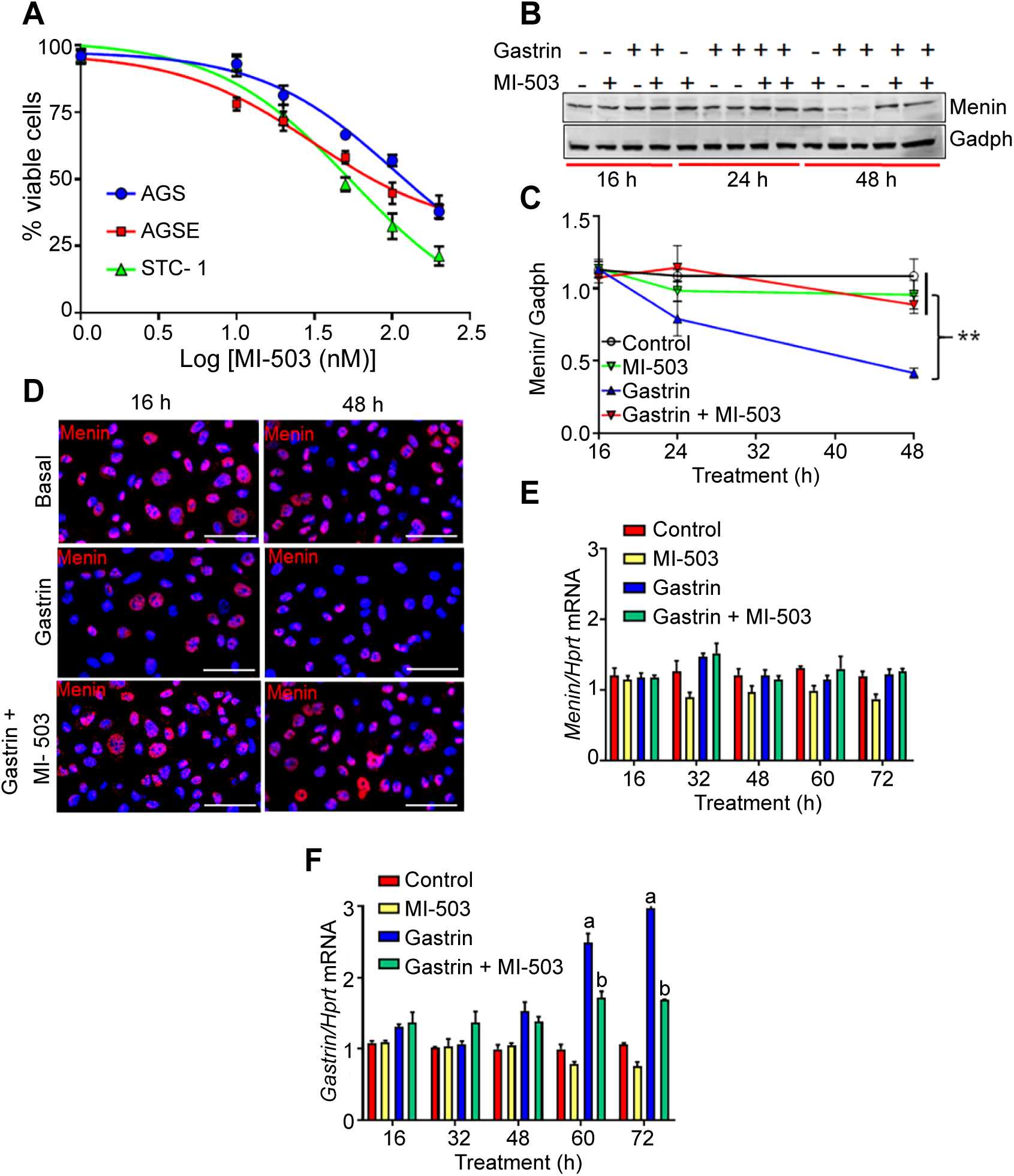
MI-503 suppresses cell growth. (**A**) Long-term viability studies using different doses of MI-503 in AGS, AGS-E, and STC-1 cells. Cell lines were treated with MI-503 for 5 days, and viable cells were normalized to DMSO controls. (**B**) Menin mRNA expression in STC-1 cells treated with or without gastrin, in the presence or absence of 50 nM MI-503 for 16-72 h. RT-qPCR for menin normalized to *Hprt* mRNA (*n* = 8 expts). (**C**) Gastrin mRNA expression in STC-1 cells treated with or without gastrin, in the presence or absence of 50 nM MI-503 for 16-72 h. RT-qPCR for gastrin normalized to *Hprt* mRNA (*n* = 8 expts). *p*<0.05 indicated by letters. (**D**) Immunofluorescent staining of menin (red) in AGS cells treated with gastrin or MI-503 for 16, 24 and 48 h. (**E**) Changes in menin were quantified by western blot and then (**F**) plotted as a function of time. Kruskal-Wallis ANOVA test with Dunn’s test of multiple comparisons was used to determine significance. Data shown are the Median ± Interquartile Range of median. ** *p<* 0.01.

To assess the impact of MI-503 gastrin gene expression in the 3 cell lines, we transfected the AGS cell line with the reporter plasmid 9.8GasLuc construct and showed that MI-503 inhibited gastrin gene expression in a dose dependent manner (**Fig. 6A**). The inactive MI-2-2 had no effect on the luciferase activity. Next, we examined endogenous gastrin mRNA in the three cell lines (**Fig. 6B**) and gastrin peptide release in STC cells (**Fig. 6C**). Both AGS and STC-1 cells generate gastrin mRNA which is suppressed with MI-503 treatment. In addition, we examined the impact of inhibiting gastrin gene expression on gastrin secretion since STC-1 cells produce amidated gastrin that can be detected by the ELISA assay. As anticipated, MI-503 inhibited gastrin peptide secretion. To examine the effect of MI-503 on the CCKBR and PKA activation, the AGS and AGSE cells were treated with MI-503. Western blot analysis showed a decrease in CCKBR levels accompanied by a strong inhibition of the catalytic PKA subunit. Taken together the results show that MI-503 inhibits PKA activity, which is a downstream effector of gastrin activation.

**Figure 6.**
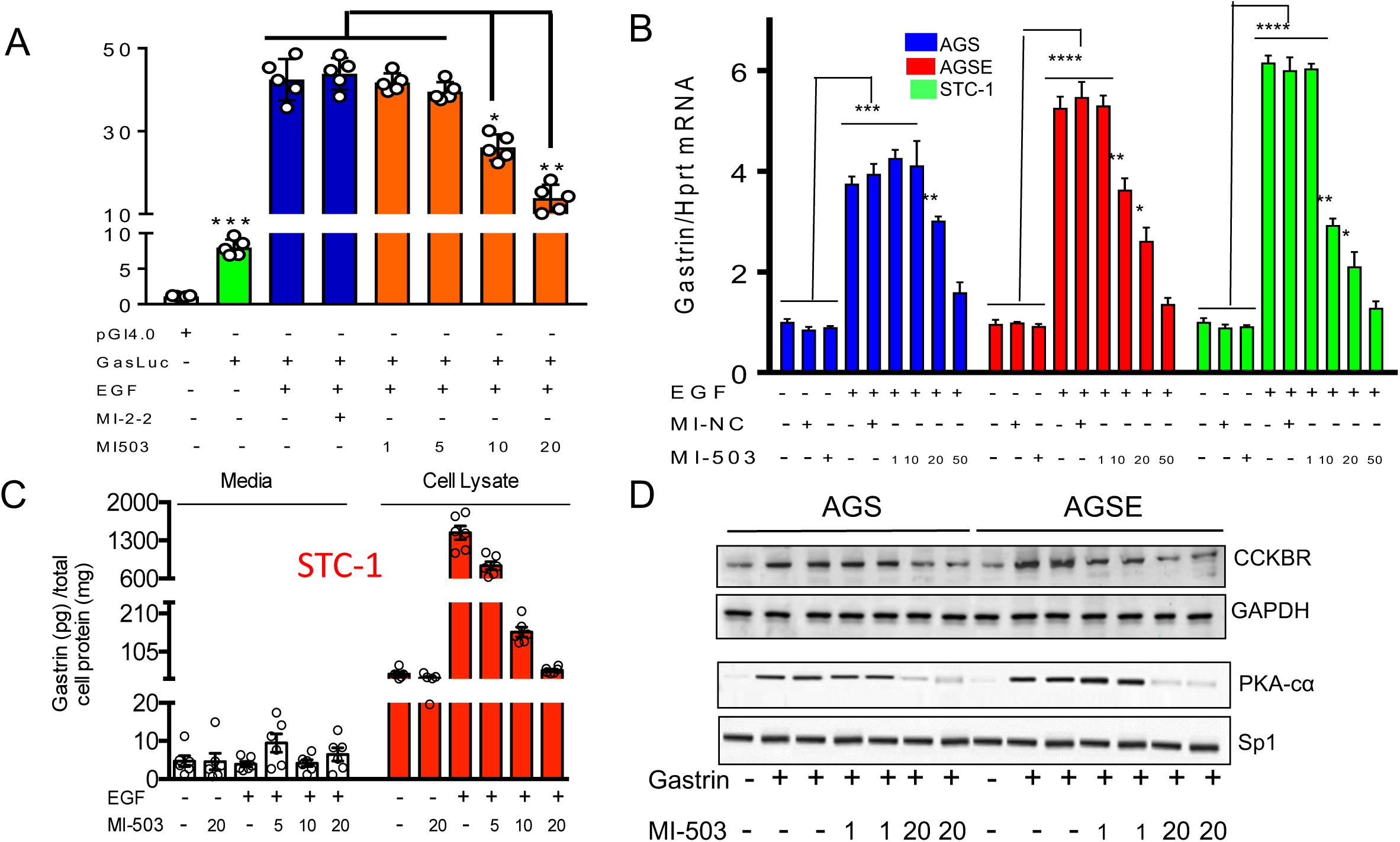
MI-503 suppresses gastrin gene expression and secretion. (**A**) MI-503 dose-dependently decreased EGF induced activation of the 9.8 kb GasLuc plasmid (pGI4.0∼9800 bp of human gastrin promoter) in AGS cells. (**B**) Similarly, EGF-induced gastrin mRNA that was reversed by 50 nM MI-503 in human AGS, AGSE, and mouse STC-1 cells. Since AGS and AGSE do not release detectable levels of gastrin in the media, we tested MI503 mediated effects on gastrin release in STC-1 cells. (**C**) EGF at 20 nM robustly increased intracellular gastrin peptide, and stimulation with 10 nM bombesin stimulated release of gastrin into the media - effects that were dose-dependently decreased by MI-503. MI-503 at 20 nM also reversed the spikes in gastrin synthesis and release. (**D**) Representative blot showing CCKBR, PKA-cα, and menin expression AGS and AGS-E cells with or without 20 nM gastrin and 1 or 20 nM MI-503.

## Discussion

In this report, we hypothesized that retaining nuclear menin would be sufficient to suppress gastrin expression and ultimately its secretion. We tested this hypothesis by treating hypergastrinemic mice with MI-503, a small molecule that binds to F9 and F13 pockets of Menin [17]. MI-503 inhibited hypergastrinemia and reduced phosphorylated PKA, menin translocation to the cytoplasm where it was ubiquitinated and subsequently degraded by the proteasome. By performing RNA-Seq on gastrin treated glial cells, we discovered that enteric glial cells express EGFR ligands epiregulin and neuregulin. Accordingly, we found that EGF receptor activation also induced Menin translocation suggesting that hypergastrinemia increases the production of EGF receptor ligands that might have an autocrine effect on glial cells. In vivo MI-503 treatment suppressed serum gastrin levels in the hypergastrinemic *OMS* mouse model due to the nuclear export of WT menin. Moreover, MI-503 also blocked EGFR-mediated translocation of Menin in vitro in both mouse and human cell lines that express gastrin. Therefore, we concluded that MI-503 suppresses gastrin in vivo and in vitro by blocking Menin translocation.

There are now a variety of mechanisms to inhibit hypergastrinemia due to gastrinomas [25, 26], but none target the genetic mutations that strongly influence de-repression of the gastrin promoter and the multiple downstream effects that follow. We have proposed that both WT Menin protein and to a greater extent mutant forms of Menin undergo translocation to the nucleus and ultimately degradation. While retaining gastrin in the nucleus may not completely suppress gastrin gene expression, it appears to dampen hormone levels. In that case, direct modulation of this nuclear factor might work in concert with other mechanisms that suppress gastrin, such as CCK2B inhibitors and somatostatin agonists (**Fig. 7**). Moreover, the identification of EGFR activation as a mechanism for stimulating menin translocation, suggests that signaling pathways downstream of growth factors likely contributes to the neuroendocrine phenotype. Targeting missense mutations of menin or those with minimal truncations might still retain their ability to suppress gene promoters and subsequent untoward effects.

**Figure 7.**
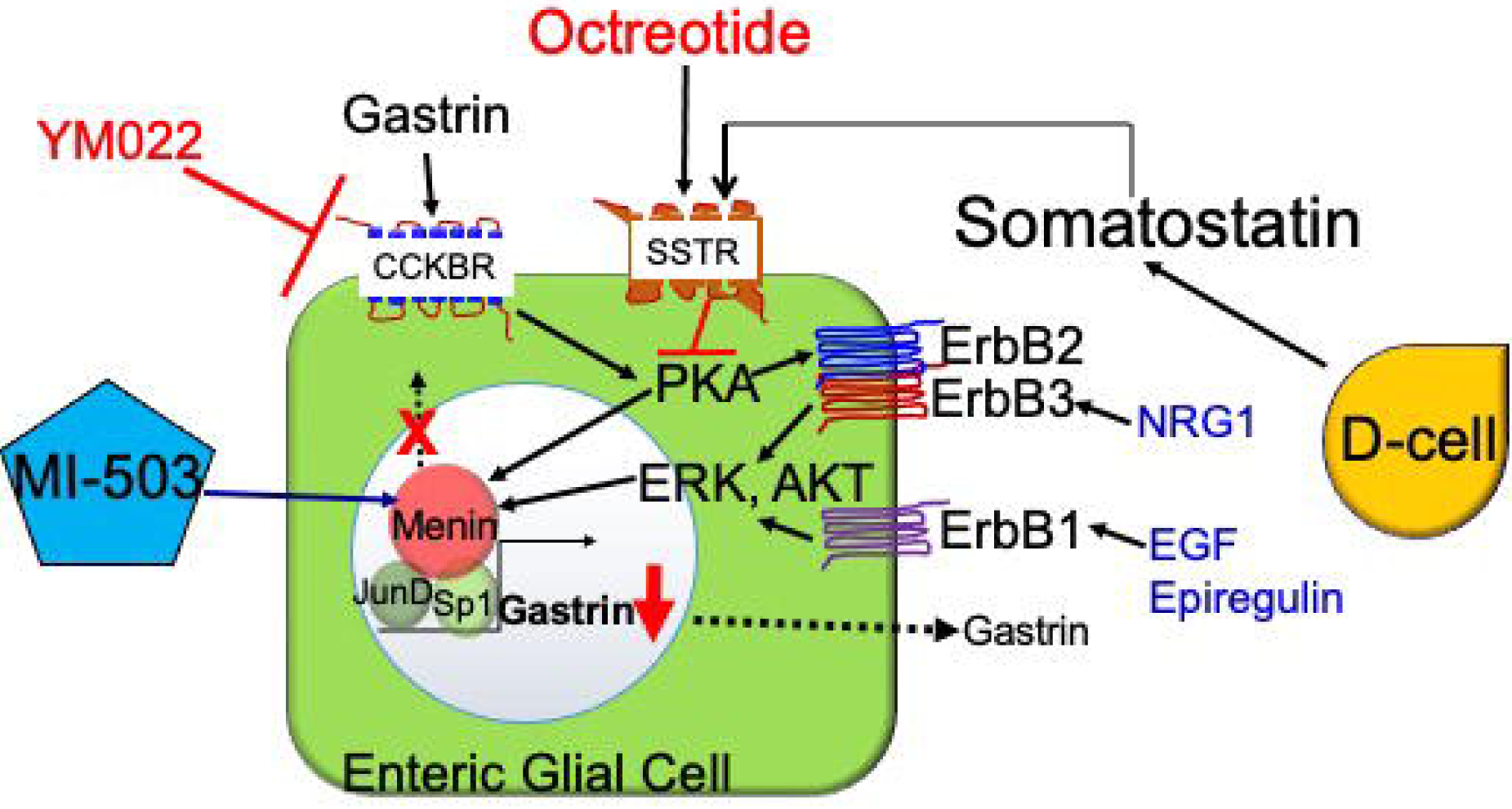
Mechanisms of gastrin repression. Here we suggest that MI-503 binds to Menin and blocks its translocation to the cytoplasm, which in turn inhibits gastrin gene expression. Other mechanisms for inhibiting gastrin include the CCKBR (gastrin receptor) inhibitor YM022 and the somatostatin receptor agonist Octreotide. We posit that activation of EGF receptor or gastrin can induce enteric glial cells to produce gastrin by inducing nuclear Menin to translocate to the cytoplasm.

The approach used in this report took advantage of a mouse model that developed elevated gastrin levels like those observed in human subjects. However, the studies in mice and human cell lines demonstrated the ability to regulate WT Menin through post-translational modification that in turn affects its cellular location. In human subjects with the MEN1 syndrome, both WT and mutant forms are present. Thus, it remains to be determined whether some *MEN1* mutations that do not correlate with the tumor phenotype, might develop tumors under specific conditions in the tumor microenvironment with signal production. Future studies will need to examine whether mutant forms of Menin are affected by MI-503.

## Methods

### Animals

All animal experiments were approved by the University of Michigan’s Committee on the Use and Care of Animals.*VC:Men1*^*FL/FL*^; *Sst*^*-/-*^(*Men1*^Δ*IEC*^; *Sst*^*-/-*^) mice were generated as previously described [6]. Mice were housed in a facility with access to food and water ad libitum. Experimental mice were fed omeprazole-laced chow (200 ppm, TestDiet, St. Louis, Missouri, USA) for 6 months to 1 year. Vehicle (25% DMSO, 25% PEG400, 50% PBS) or MI-503 (10, 30, and 55 mg/kg body weight) was administrated intraperitoneally, once daily for 4 weeks. Blood was drawn via submandibular bleeding prior to MI-503 administration to determine basal plasma gastrin levels. At the end of MI-503 treatment, blood was drawn from the mice to determine effects of the drug. Mouse genders used were equivalently distributed across the control and treatment groups.

### Plasmids

The human gastrin luciferase vector containing 9800 bp of the human gastrin promoter was previously described [22].

### Cell lines

Human gastric adenocarcinoma cell line (AGS) cells and STC-1 cells derived from a mouse intestinal tumor [27-29] were purchased from ATCC. AGS cells stably expressing the CCKBR (AGS-E) were a gift from Dr. Timothy Wang, Columbia University [30]. Both AGS and STC-1 cells were cultured in Dulbecco’s modified Eagle’s medium (DMEM; Invitrogen, Carlsbad, California, USA) supplemented with 10% fetal bovine serum (FBS).

### Primary Glial Culture Isolation

The first 6-8 cm of the proximal duodenum from 2 mice were used for the isolation of duodenal enteric glia and pooled as described previously [5]. Briefly, the longitudinal muscle/myenteric plexus (LMMP) attached to the submucosa was removed from the underlying circular muscle by gently rubbing the edges of forceps and teasing it away using a cotton swab wetted with DPBS by applying light horizontal strokes. Next, the epithelium was removed by incubating the tissue twice in EDTA/HEPES/DPBS (5mM/10 mM) solution at 4°C for 10 min with gentle rocking. The tissue was gently titurated 5 times using a 5 ml pipette to dislodge the epithelium. The tissue was recovered on a 100µm filter. After the last incubation, epithelial-stripped tissue was incubated in Cell Recovery Solution® (Corning) for 30 min at 4°C with rocking then titurated 3 times to generate the glial cell containing solution, filtered through a 40µm filter to retain the tissue and large clumps. The cell suspension that flows through was spun at 2000 x g for 5 min at 4°C and then re-suspended in 1.2 mL of DMEM/F12 medium supplemented with 10% FBS, 100 IU/mL penicillin, 100 µg/mL streptomycin, 20 µg/mL gentamicin, 6mM glutamine, and 2.1 g/L NaHCO_3_. About 100-200 µL of dissociated cell suspension was transferred to laminin and poly-D-lysine coated 6-well plate containing DMEM/F12 medium supplemented with B-27, 2 mM L-glutamine, 1% FBS, 10 µg/mL Glial Derived Neurotrophic Factor (GDNF), 100 IU/mL penicillin, 100 µg/mL streptomycin, and 20 µg/mL gentamicin. The cells attach by 16 hours. After changing the media within 24h of attachment, the cells remain viable for 4 days.

### RNA-Seq and Validation

Primary enteric glial cells were prepared from the proximal duodenum and cultured for 3 days prior to treating with 20 nM gastrin or omeprazole for 8 h. The cells were then removed from the plate and submitted for bulk RNA-Sequencing (University of Michigan Sequencing Core). Sequence data was deposited into iPathways v1.2 (Advaita Corp) for analysis. Recombinant mouse Epiregulin was purchased from Sino Biological and mouse neuregulin1 from R&D Systems.

### RNA extraction and RT-qPCR

Total RNA was isolated from tissues, glial cultures, and cell lines using the RNeasy Mini kit (Qiagen). About 500 ng of total RNA was used for cDNA synthesis (iScript cDNA synthesis kit, Bio-Rad, Hercules, California, USA). cDNA was diluted 1:5 before use in PCR reactions. Quantitative PCR (qPCRs) was carried out using a thermal cycler (model C1000, Bio-Rad) with Platinum Taq DNA polymerase (Invitrogen) and SYBR Green dye (Molecular Probes, Carlsbad, California, USA). PCR reactions were performed in duplicate using the following conditions. Gastrin (mouse and human): 95°C for 1 min, 39 cycles of 95°C for 9 s and 65°C for 1 min followed by 55°C for 1 min. CCKBR - 95°C for 1 min, 39 cycles of 95°C for 9 s and 62°C for 1 min followed by 55°C for 1 min. Menin (mouse and human): 95°C for 1 min, 39 cycles of 95°C for 9 s and 60°C for 1 min followed by 55°C for 1 min. The primers used are listed in Supplemental Table S1. Differences in mRNA expressions were normalized to *Hprt* and then expressed as fold increase over indicated controls.

### Western blot

Total cellular proteins were extracted by homogenization in radio-immunoprecipitation assay (RIPA) lysis buffer (Sigma-Aldrich) supplemented with the Complete Protease Inhibitor Cocktail (Roche, Indianapolis, Indiana, USA) and PhosSTOP phosphatase inhibitor cocktail (Roche). Nuclear and cytoplasmic fractions were isolated using the NE-PER kit (Thermo Scientific), as per manufacturer’s instructions. For enteric glial cell experiments were performed by hypotonic lysis in 200 µl of buffer A (20 mM HEPES, pH 8.0, 1 mM DTT, 0.1% NP-40 and a proteinase inhibitor cocktail from Roche). The cell lysates were incubated on ice for 10 minutes, and then centrifuged at 400g for 5 min at 4C. The supernatants were designated as the cytoplasmic fractions. The resulting pellet containing the nuclei were washed 3 times in buffer A and then lysed in 50 µl of buffer B (20 mM HEPES, pH 8.0, 20% glycerol, 500 mM NaCl, 1.5 mM MgCl2, 0.2 mM EDTA, PH 8.0, 1 mM DTT, 0.1% NP-40 and a proteinase inhibitor cocktail) for 30 min on ice. The buffer B suspension was centrifuged at 15,000g at 4C for 15 min. Cellular lysates were resolved on Novex 4%– 20% Tris-Glycine precast gels (Life Technologies) and resolved proteins were transferred to polyvinylidene difluoride (PVDF) membranes. Membranes were blocked in 5% BSA to prevent non-specific binding and incubated with primary antibodies overnight at 4°C (see online supplementary table S3). Following incubation (1 hour, RT) with infrared dye-labeled secondary antibodies, IRDye 800CW and IRDye 680RD (LI-COR), target proteins were detected using the Odyssey Infrared System (LI-COR Biosciences, Lincoln, NE). Bands were quantitated using the LI-COR Odyssey software, version.

### Immunohistochemistry

Duodenums of mice were flushed with ice-cold PBS, cut open and fixed overnight in 10% formaldehyde solution at 4C. They were then transferred to 70% ethanol and processed before embedding in paraffin (TissueTek). Five micron sections were cut from paraffin blocks for staining. Sections were deparaffinized in xylene and dehydrated by brief sequential incubations in 70%, 90%, and 100% ethanol. Antigen retrieval was performed by warming the slides in Tris-EDTA (pH 9.0), or sodium citrate (pH 6.0) for 30 min. Non-specific binding was blocked by incubation in 10% donkey serum followed by incubation with primary antibodies (2 h at RT, or overnight at 4°C, see online Supplementary Table 1, list of antibodies). For fluorescent detection, sections were incubated (RT) with Alexa Fluor-conjugated secondary antibodies for 30 min (1:400 dilution). ProLong Gold antifade reagent with 4′, 6-diamidino-2-phenylindole (DAPI, Invitrogen) was used for nuclear counterstaining. Images were taken using a Nikon inverted confocal microscope (Nikon, New York, USA).

Primary glial cultures were plated on laminin and poly-D-lysine coated coverslips, fixed using 4% paraformaldehyde for 20 min (RT), and then permeabilized for 5 min with 0.2% Triton X-100/phosphate buffered saline (PBS). Cultures were then blocked in 1% serum (from the appropriate species) for 30 min (RT), incubated with primary antibodies, and stained as described above.

### Measurement of Gastrin in plasma, tissues, and primary cultures

For plasma gastrin, mice were fasted for 16h before euthanization. Blood was collected by cardiac puncture in heparin-coated tubes and plasma was collected by centrifugation at 5000g for 10 min at 4°C. About 50 µL of the plasma was used for measuring gastrin levels using the Human/Mouse/Rat Gastrin-I Enzyme Immunoassay Kit (RayBiotech, Georgia, USA), per the manufacturer’s instructions. For measuring gastrin content of duodenal or antral tissue, 10-20 mg of antral and duodenal tissues were boiled in 300 µL of deionized water. The extract was spun briefly at 2000g and the supernatant was used for gastrin analysis as described above. Gastrin content was expressed after normalization to tissue weight. For measuring gastrin content of primary glial cultures, cultures were scraped in PBS using cell scraper. Cells were pelleted, and then boiled in deionized water (volume equals to 10 times the size of the pellet). The extract was spun briefly as described above and supernatant was used for gastrin measurements. Gastrin content was expressed after normalization to total protein content.

Gastrin secreted from primary cultures in response to 10 nM bombesin was also measured. Three to four day old primary glial cultures were serum-starved for 4 hours, and stimulated for the indicated times. Media collected was concentrated using protein concentrators PES, 10K MWCO (Thermo Fisher Scientific). About 2 mL of media was concentrated to ∼ 170 µL by centrifuging in a swinging-bucket rotor at 4000g for 60 min at 25°C. ELISA was used to determine the amount of gastrin peptide in the concentrated media and tissue extract by after normalizing to protein content.

### Ubiquitination Assay

Ubiquitinated proteins were pulled down using the Agarose-Tandem Ubiquitin Binding Entities (Agarose-TUBE, Life sensors). Cells were lysed in cell lysis buffer containing 50mM Tris-HCl, pH 7.5, 0.15M NaCl, 1mM EDTA, 10 mM NEM, 1% NP-40, and 10% glycerol supplemented with protease and phosphatase inhibitor cocktail. About 20µl of resin in 500µl of lysis buffer containing 1-2mg of total protein was used per reaction. Clarified cell lysates were incubated with uncoupled agarose (Life sensors) for 30 min at 4°C with rocking; agarose was removed by centrifugation and the clarified supernatant was used as control for non-specific binding. An aliquot of each sample was removed prior to immunoprecipitation and designated “INPUT”. The remnant cell lysates were incubated with equilibrated Agarose-TUBEs overnight at 4°C on a rocker. Beads were collected by low speed centrifugation (3000g, 4°C) for 5 minutes, washed with TBS-T. Bound protein was eluted by boiling for 5 min in Laemmli sample buffer.

### MTT Assay

MTT assay was performed using the Cell Proliferation Kit (Sigma Aldrich) as per manufacturer’s instructions. Briefly, about 1000 AGS, AGS-E, and 2000 STC-1 cells were grown in 96-well plates in a final volume of 100µL of culture medium. After 96 hours of treatment with MI-503 or vehicle, 10µL of the MTT labeling reagent (0.5 mg/mL) was added and incubated for 4 h at 37°C. Then, 100 µL of the solubilization reagent was added and incubated overnight at 37°C. To ensure complete solubilization of the purple formazan crystals the plate was shaken for 1 h and absorbance was read at 560 nm, with reference wavelength of 660 nm. Cell Viability (%) was calculated as: {OD_560_ – OD_660_ of treated group/OD_560_ – OD_660_ of control group} * 100.

### Statistical analyses

All reported values are the mean ± SEM. Statistical analyses of data obtained from mice tissues and primary cultures were performed using the non-parametric Kruskal–Wallis test (GraphPad Prism 6). Dunn’s multiple comparison test was used to identify groups that were significantly different. Data from *in vitro* sample sets was analyzed using one-way analysis of variance (ANOVA), followed by Tukey’s multiple comparisons test. P<0.05 was considered significant.

## Abbreviations

*Men1*^*ΔIEC*^: *VillinCre;Men1*^*FL/FL*^
*Sst*^*-/-*^: somatostatin null
*OMS*: <u>O</u>meprazole-treated <u>M</u>en1^ΔIEC^;<u>S</u>st^-/-^
GFAP: glial fibrillary acidic protein
*GAST*: gastrin

## Figure Legends

**Supplementary Table 1:** Antibody Sources.

| Antibodies | Vendor (Cat#) | Dilution/Application | Antigen Retrieval |
| --- | --- | --- | --- |
| Gastrin | Dako (A0568)<br>Santa Cruz<br>Biotechnology (sc-7783) | 1:1000 (tissue), 1:200 (cells)<br>1:400 (tissue) | Tris-EDTA, pH 9 |
| Menin | Bethyl Laboratories<br>(A300-105A) | 1:10,000 (tissue with TSA,<br>mouse sections)<br>1:2000 (cells with TSA,<br>human sections)<br>1:500 (cells)<br>1:1000 (WB)<br>2 µg/mg of lysate (IP) | Sodium citrate |
| GAPDH | Cell Signaling<br>Technology (2118) | 1:3000 (WB) | --- |
| β-actin | Santa Cruz<br>Biotechnology (sc-130656) | 1:3000 (WB) | --- |
| Sp1 | Santa Cruz Biotech<br>(sc-14027) | 1:2000 (WB) | --- |
| Ubiquitin | Abcam (ab7254) | 1:100 (WB)<br>1:400 (cells) | --- |

WB: Western Blot; IP: immunoprecipitation; TSA: Tyramide Signal Amplification

